# Avoidance Of Rejuvenation: A Stress Test For Evolutionary Theories Of Aging

**DOI:** 10.1101/2025.08.24.671987

**Authors:** Samir Aisin, Boris V. Lidskii, Peter V Lidsky

## Abstract

The biological feasibility of human rejuvenation remains a subject of intense debate, yet answering this question is critical for guiding research strategies. Should aging research focus on reversing aging in older individuals, or on pausing its progression at earlier ages? We address this question with evolutionary biology.

Classic evolutionary theories of aging— damage accumulation, antagonistic pleiotropy, and the disposable soma—consider aging as a detrimental byproduct of evolution. From this perspective, rejuvenation should confer strong fitness advantages and therefore be expected to evolve in species experiencing substantial aging in the wild. Its rarity in nature should thus be interpreted as evidence of its mechanistic implausibility.

Yet, rejuvenation does occur in a few species, and, paradoxically, it is typically induced by stress but not used under optimal conditions. Using mathematical modeling of lifespan plasticity in eusocial insects, we show that this pattern cannot be reconciled with classic theories of aging, revealing an internal contradiction between these theories and the observed avoidance of rejuvenation. By contrast, the pathogen control hypothesis—which interprets aging as an adaptive, programmed process—offers a consistent evolutionary framework for understanding and potentially achieving rejuvenation.

## Introduction

Physiological rejuvenation is humanity’s most ambitious aspiration and the ultimate goal of biogerontological research. Recent advances in partial cellular reprogramming and regenerative biology have fueled optimism that physiological rejuvenation may be achievable. Yet whether rejuvenation is fundamentally possible remains deeply contested. Some authors consider systemic rejuvenation within the long-term reach (Zhang et al., 2022), while others claim it is theoretically impossible (Tarkhov et al., 2024). This debate is complicated by the absence of universally accepted definitions of “aging” and “biological age,” and by widely diverging opinions on aging’s primary causes and mechanisms (Gladyshev et al., 2024). However, understanding the feasibility of rejuvenation is critical for informing experimental research: should efforts focus on rejuvenating old organisms or slowing aging progression in middle-aged individuals?

To evaluate the feasibility of physiological rejuvenation, we use evolutionary approaches. First, we assess the known examples of rejuvenation, analyze the ecological context that may trigger animals to rejuvenate, and use mathematical modeling to reconcile these observations with different evolutionary hypotheses. Our goal is to resolve a central paradox: if rejuvenation is mechanistically possible, why has evolution not widely adopted it as a strategy to extend lifespan and enhance fitness?

### Definitions

Living organisms possess an intrinsic characteristic, known as *biological age*, that gradually increases over time through a process called *aging*. Aging has a negative impact on an organism’s health, ultimately leading to its death. Changing the tempo of aging in response to the intrinsic or extrinsic stimuli is termed *aging plasticity*. Plasticity includes a faster increase in biological age, or *accelerated aging*, e.g., in response to inflammation(Franceschi et al., 2018; Furman et al., 2019) or mating (Chapman et al., 1995), and, oppositely, slower or *decelerated aging*, e.g., in response to dietary restriction(Nakagawa et al., 2012). *Rejuvenation* is a specific type of aging plasticity manifested in decreasing biological age over time. Rejuvenation can be *comprehensive*, meaning that all physiological parameters are reverted, and the organism becomes indistinguishable from its younger self. Alternatively, rejuvenation can be *partial*, meaning that only some age-associated physiological parameters are reversed. Evidence for rejuvenation can be divided into two types: *reverse development*, when an animal metamorphoses into an earlier developmental stage, and *adult rejuvenation*, when the functions of an adult individual revert to a more youthful state.

### Examples of aging plasticity and physiological rejuvenation

Rejuvenation ubiquitiously occurs during development, e.g., in mammals in the gastrulation stage (Kerepesi & Gladyshev, 2023). Thus, the question of the feasibility of rejuvenation is now limited to the later developmental stages.

The immortal jellyfish (*Turritopsis dohrnii*) is the most renowned example of reverse development in animals. The typical jellyfish lifecycle includes an attached polyp stage that later transforms into a free-swimming medusoid that reproduces and dies. It was shown that under starvation, suboptimal temperature, or chemical stress, medusoids can revert into polyps and escape death this way(Piraino et al., 1996). Recently, a similar ability to reverse development in response to stress was discovered in the ctenophore, *Mnemiopsis leidyi* (Soto-Angel & Burkhardt, 2024). Observations made in these very distant species indicate the conserved pattern: although rejuvenation is mechanistically feasible, paradoxically, animals resort to it only when under stress.

Rarely, reverse development can be found outside the context of stress. In some termite species belonging to the *Archotermopsidae* and *Kalotermitidae* families, later larvae can undergo reverse development into earlier larvae under optimal conditions(Korb et al., 2021; Korb & Hartfelder, 2008).

Evidence for rejuvenation without metamorphosis is scarce. In vertebrates, the regeneration of limbs in axolotls results in decreased tissue biological age as judged by methylation clocks (Haluza et al., 2024). Additional evidence supporting the feasibility of tissue rejuvenation comes from laboratory experiments using Yamanaka factor-mediated partial reprogramming (Chondronasiou et al., 2022; de Lázaro et al., 2023) and perfusions with young plasma (Zhang et al., 2023) in mice and rats. These interventions result in the reversion of multiple physiological parameters to their more youthful states. Altogether, these results support the feasibility of at least partial physiological rejuvenation in mammals.

While encouraging, this conclusion poses a fundamental paradox: if rejuvenation is mechanistically possible, why didn’t organisms evolve to activate it every time they reach old age and prefer dying instead? Why did evolution not employ rejuvenation to increase older organisms’ fitness? Since evolution universally probes all available opportunities, the rarity of rejuvenation in nature must indicate its mechanistic impossibility. However, the abovementioned examples of hydrozoans, ctenophores, and termites fail to fit this general model. Resolving this contradiction is critical for instructing future research and developing rejuvenating therapies. Moreover, this may help us test and refine the existing theories of aging and assist in the search for the unifying paradigm, currently missing from the field (Gladyshev et al., 2024).

Eusociality, a colonial social structure characterized by the division of labor between breeder and non-breeder individuals, is also associated with the most pronounced examples of aging plasticity, which may or may not involve metamorphosis. Different castes have different aging rates: breeders age more slowly than non-breeders in most species of eusocial insects and mole-rats (families *Bathyergidae* and *Heterocephalidae*). In mole-rats, a non-breeder can undergo reproductive activation and become a breeder. In this case, it ages at a slower pace (Dammann et al., 2011; Horvath et al., 2022). Similar phenotypes are observed in some ant species: if the queen dies, several workers in the colony may undergo reproductive activation and become “gamergates,” pseudo-queens that can live approximately five times longer than unperturbed sterile workers. Gamergates can also revert to a worker caste and go back to faster aging(Ghaninia et al., 2017; Yan et al., 2022). These essential results imply the existence of an aging master regulator that can orchestrate all the processes that define changes in biological age simultaneously. However, the existence of an aging master regulator does not prove rejuvenation. These phenotypes can be rationalized by pausing aging, a state in which biological age increments very slowly but does not reverse.

An example, mostly resembling physiological rejuvenation in adults, can be observed in honeybees (*Apis mellifera*). After leaving the pupae, the honeybee is committed to working as a nurse, cleaning the nest and feeding larvae and the queen. Nurses do not leave the colony. After some weeks, nurses transition to foraging and begin collecting nectar and pollen outside the hive. This job transition is associated with a dramatic shift in the honeybee physiology: the protein storages decrease, specifically the amounts of a regulatory and storage protein vitellogenin; the levels of juvenile hormone increase, inducing the immune cells (hemocytes) to die via pycnosis (Amdam et al., 2005). Foragers work for a few more weeks, deteriorate, and die because of aging or extrinsic reasons. However, if the number of nurses declines as a result of a predator attack or experimental manipulation, some of the foragers must return to the nursing job. In this case, they undergo a physiological rejuvenation: protein storage and vitellogenin levels increase, the number of hemocytes is restored, and the mortality decreases. Transcriptomics and methylomics profiles also revert to a more youthful, nursing stage(Guan et al., 2013; Margotta et al., 2012).

Lifespan extension can also be observed in honeybee laying workers. In the case of the queen’s death, workers undergo reproductive activation and become long-lived fertile isoforms (Paleolog et al., 2021), similar to gamergates in ants.

Although it remains unclear if honeybee rejuvenation is comprehensive or partial, it is evident that adult animals have some potential for reverting physiological decline and extending lifespan. Nevertheless, as in cases with jellyfish and ctenophores, honeybees and most other eusocial animals resort to rejuvenation or lifespan extension only under sub-optimal conditions, e.g., if queens or nurses die. But why don’t they activate their lifespan-extending potential if not under stress? One of the traditional takes is that foragers underinvest in their maintenance due to high extrinsic mortality. If an animal will not survive to a late age, underinvestment might be a good energy-saving strategy. However, regardless of high extrinsic mortality, aging was shown to account for about 7% of foragers’ deaths (R. Dukas, 2008). Hence, the activation of lifespan-prolonging mechanisms during the normal lifecycle must save a fraction of foragers and therefore be evolutionarily beneficial, as it would save energy and increase the hive’s fitness. Why aren’t the existing lifespan plasticity mechanisms activated in these cases?

To answer this question, we use mathematical modeling to rationalize this observation in the context of different theories of aging.

We found that while classic models – damage accumulation (Gladyshev, 2016; Lopez-Otin et al., 2023; Ogrodnik et al., 2019), antagonistic pleiotropy (Williams, 1957), and disposable soma (Kirkwood & Holliday, 1979) – fail to explain phenomena and are incompatible with the existence of rejuvenation and pausing aging, the pathogen control hypothesis (Lidsky & Andino, 2020, 2022; Lidsky et al., 2022) can nicely fit the data and reconcile it with evolutionary theory.

## Results

### Modeling the honeybee colony

We have modeled a honeybee community that includes *N* foragers. Foragers collect food, specifically pollen, that is considered a universal nutritional equivalent in this model. For simplicity, we did not account for proteins and carbohydrates separately. The yield of a single forager *S* is assumed to decrease with the colony’s growth since foragers need to spend more time reaching the feeding location. We calculated the yield of the n^th^ forager according to the formula:

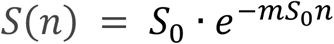

where *S*_0_ is the yield of the very first forager, and 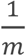 is the maximum possible yield of the hive for any number of foragers. The cumulative yield of a hive consisting of n foragers is assumed to be:

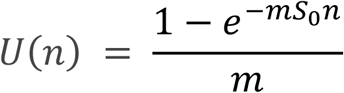

We studied colonies that have optimal amounts of foragers, meaning that the number of foragers is *N*, and 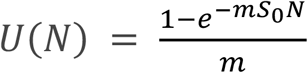.

Forager’s productivity was assumed not to decline with age, as it was demonstrated at least within the first seven days of foraging (Reuven Dukas, 2008).

The colony uses collected pollen in the following ways: (i) to produce new foragers, (ii) to feed and maintain the existing foragers, and (iii) to produce new queens that will establish new colonies.

i. Feeding foragers. The pollen a forager consumes daily (*M* + *F*) is utilized in two ways: the feeding component *F* goes towards supporting physical and physiological functions, such as flight, digestion, and cognition. Maintenance amount *M* is invested in repairing physiological damages *D* that each forager receives daily and that classic theories assume to be the primary causes of aging (Lopez-Otin et al., 2023). The difference between *M* and *D* defines the maximum lifespan: if the damages are not repaired completely (*M* < *D*), they accumulate and kill the forager when reaching the value of *C* ∗ *L. C* is the nutritional cost of a forager expressed in mg of pollen. *L* is the proportion of the body’s cells and molecules that, if damaged, kill the animal (0 < *L* < 1). Therefore, if *M* < *D*, the forager will die of accumulated damage in

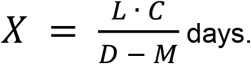 In addition to the death of old age, foragers are exposed to age-independent extrinsic mortality with a probability of death *μ* per day.
ii. Production of new foragers: *n*_0_ are made per day. In the steady-state colony

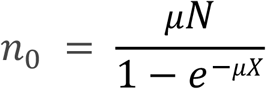

and the total amount of energy invested in the production of new foragers is therefore *C* · *n*_0_ or 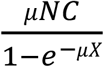.
iii. The rest of the pollen is invested in the production and maintenance of nurses and queens and defines the reproductive fitness of the hive. This remaining value is calculated as:

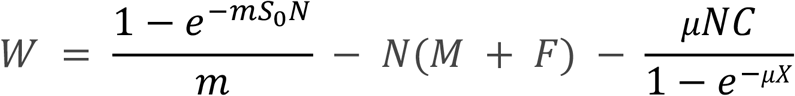

Assuming the colony invests its resources to maximize reproductive fitness, we sought to determine how *W* depends on *n*0 and *M*. Initially, we assumed no lifespan plasticity (*M* is constant) as described (Fig. 1; Table I) (Kramer & Schaible, 2013).

**Table I.** Model parameters.

| Variab<br>les | Meaning | Values (range) | Units | Comment |
| --- | --- | --- | --- | --- |
| N | Number of foragers in the hive | variable | units |  |
| $\mu$ | Extrinsic mortality (probability of dying per day, independent of aging) | 0.13 or variable ($0 < \mu < 1$) | fraction | (R. Dukas, 2008) |
| C | Cost to produce a new forager | 137 | mg of pollen |  |
| S(n) | Yield of a single n<sup>th</sup> forager per day | $S_0(1 - m \cdot U(n))$ | mg of pollen | |
| S<sub>0</sub> | Yield of the very first forager in the hive | 230.25 | mg of pollen | Calculated with the average production per forager per day of 89.7 (Layek et al., 2020) and $\frac{1}{m} = 10^6$ |
| U(n) | Daily yield of the hive consisting of n foragers | variable | mg of pollen | $\sum_{i=0}^N S_i$ |
| 1/m | Maximum yield of all forages with any N | $10^6$ | mg of pollen | Approximation based on the estimated amount of ~11 thousand foragers per nest |
| F | Cost of feeding a single forager per day | 3.85 –M | mg of pollen | $F + M = 3.85$ (Brodschneider & Crailsheim, 2010) |
| M | Investment in the repair of a single forager per day | variable | mg of pollen |  |
| D | Daily molecular damage to a single worker | variable | mg of pollen | Measured in the amount of pollen necessary for repair |
| L | The proportion of the body that, if damaged, results in death | variable | fraction |  |
| X | Age of death due to old age | variable | days | $X = \frac{L \cdot C}{(D - M)}$ |
| W | Darwinian fitness of the colony | variable | mg / day |  |
| $n_0, n_1, \dots, n_j$ | The age structure of the bee population. $n_0$ is the number of newborn foragers, $n_1$ - one-day-olds, etc. | variable | units | $N = \sum_0^{\infty} n$ |
| N<sub>s</sub>, N<sub>i</sub> | Number of uninfected and infected foragers | variable | units | Only in Model III |
| $\beta$ | Pathogen transmission rate | variable | efficacy | Number of hosts infected by one infected individual per day in a densely packed population of healthy individuals. Only in Model III |

**Fig. 1.**
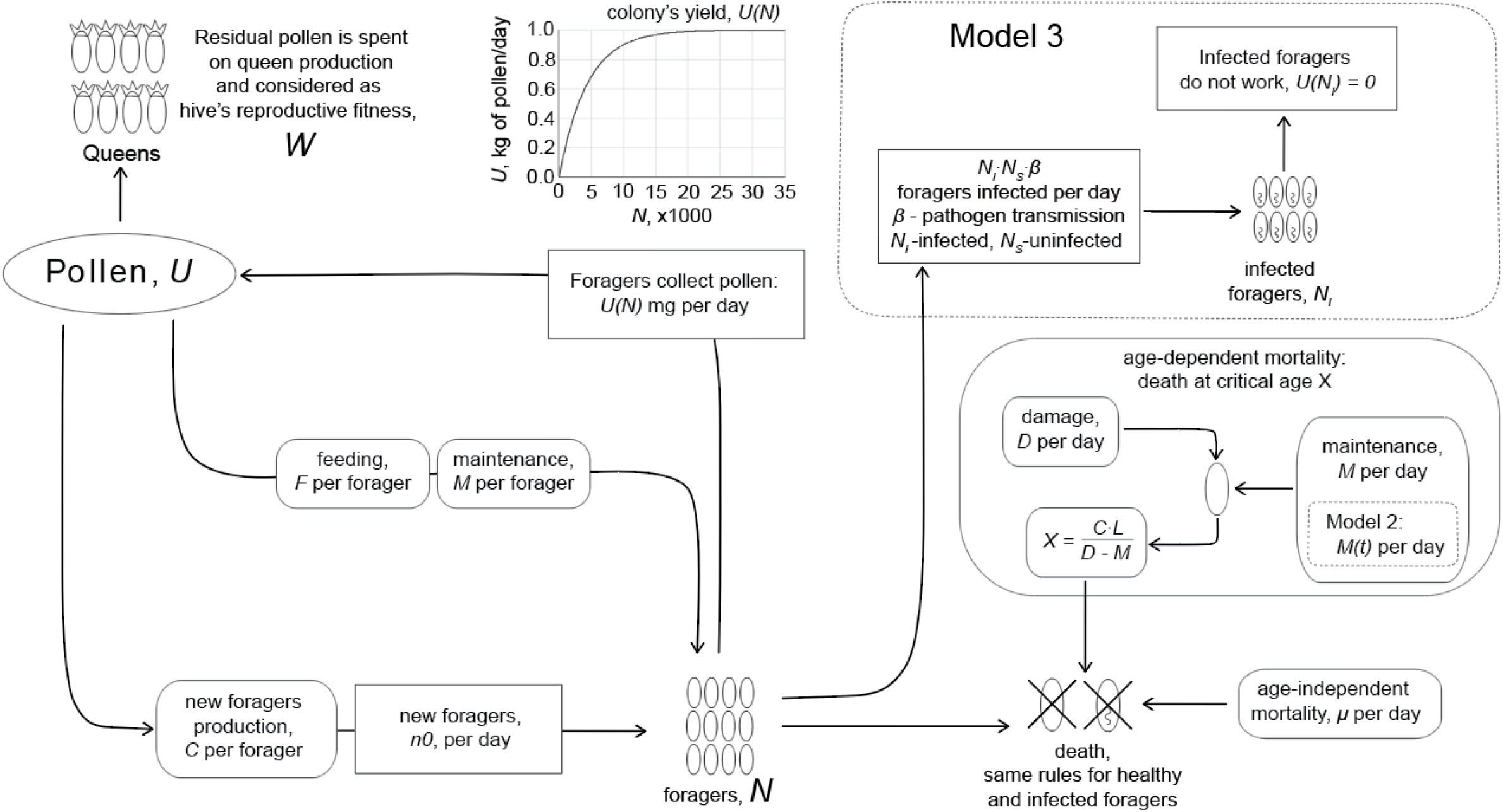
Model architecture.

### Model I

The key assumption of this model is that the investment in repair *M* is stable at all ages and cannot change. If only a few foragers may survive into the late ages due to extrinsic mortality, the costs of replacing these with new foragers are small, and such are the energy losses caused by aging to the hive. However, younger foragers comprise the majority of the colony, and the maintenance of these animals incurs substantial costs. The optimal investment strategy is found when producing additional foragers to replace those who died of aging becomes cheaper than increasing investments into better maintenance of younger cohorts.

We found that there is an optimal finite lifespan of a forager that depends on L and μ according to the formula:

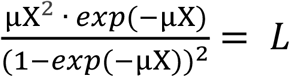

(See **Supplementary Material I** for details). Thus, if *M* is constant, our model can explain aging in honeybees, a small proportion of foragers may indeed die of age-related reasons (Fig. 2).

**Fig. 2.**
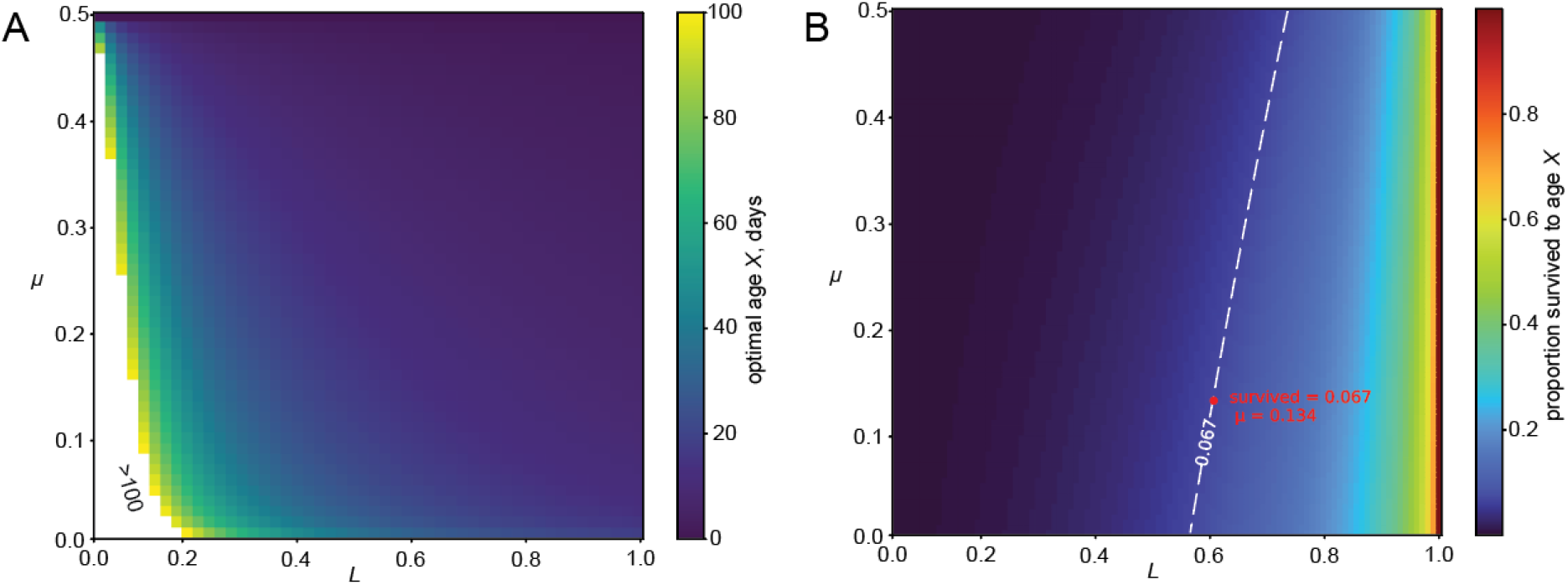
Model I outcome. **A**. The ecologically optimal *X* age of death due to aging, under the assumption that investment in maintenance, *M* is constant and evolved to maximize colony fitness. The age has been calculated for different values of extrinsic mortality μ, and the critical level of damaged tissues, *L*. **B**. The proportion of foragers that die due to aging. The red dot marks the combination of parameters when the proportion of survivors to the age of death is 0.067 and the extrinsic mortality is 0.013, as observed in (R. Dukas, 2008).

Similar results were demonstrated before (Kramer & Schaible, 2013).

### Model II

Next, we introduced the concept of aging plasticity, modeled as M, which can vary with age *t* (Fig. 1B). In this case, the optimal age-dependent investment strategy evolved to maximize colony fitness, *W* will be represented by a function *M(t)*:

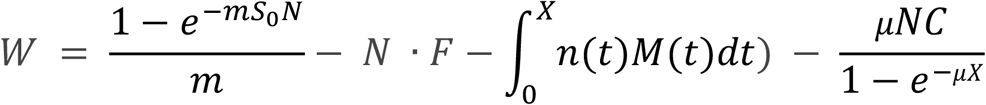

Evolution should optimize M(t) in a way that 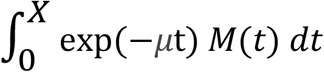 is minimal. Suppose the age of death if no investments in repair are made (*M* = 0), *X*_0_ = *LC* / *D*

Let’s examine a strategy involving no investment in repair at a young age until animals almost reach the age of death. At this point, they increase the investment up to the value of *D* which allows to stop aging and persist indefinitely:

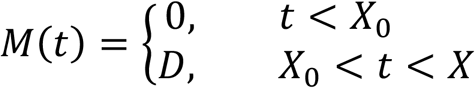

Then,

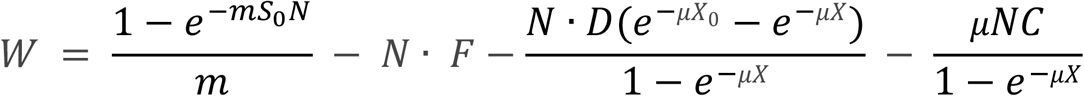

Since in the steady state situation 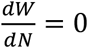,

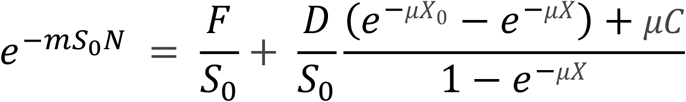

By designating the right part of this equation as η and solving

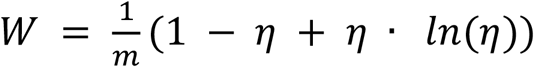

we found that this investment strategy is optimal independently of any model parameters **(Supplementary Material II)**. At the same time, we found that always 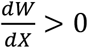, hence, a longer forager lifespan always benefits the hive, and any opportunity to extend their lifespans should be used.

Therefore, if the model allows the investment in maintenance to vary with time, young honeybees would underinvest in their maintenance to conserve energy when in large numbers. Later, the few individuals lucky enough to survive to older ages will allocate more resources to repairing damage and preventing deaths. According to this calculation, death by aging should never be observed in honeybees or any other species with existing mechanisms of aging plasticity. If energy is the currency that defines life history, and if a mechanism for prolonging lifespan exists, such a mechanism should be activated every time an individual reaches old age. This prediction contradicts the empirical data: aging is observed in honeybees (R. Dukas, 2008), while cases of increased investment in repair and longevity in older organisms have never been demonstrated to the best of our knowledge.

This disconnection indicates a fundamental incompatibility between the classic evolutionary theories of aging and lifespan plasticity. Critically, our results demonstrate that not only rejuvenation but also the pausing of aging cannot fit these theories. While physiological rejuvenation is a rare phenotype found only in a few species, pausing aging of different types is very widespread in animals, which significantly expands the ecological significance of our findings(Wilsterman et al., 2021). As such, a different theory is needed to explain the phenomena.

### Model III

Next, we tested whether evolutionary avoidance of prolonged lifespan and rejuvenation can be rationalized by a programmed aging model, specifically the pathogen control hypothesis. According to this model, aging evolved not as a side effect of underinvestment in repair, but as an adaptive program that limits the transmission of chronic infectious diseases (Lidsky & Andino, 2020).

To test this opportunity, we introduced a chronic pathogen that can spread between bees and makes foragers stop working. Such effects of the parasites were found in ants (Beros et al., 2021) and eusocial wasps (Beani et al., 2021). For simplicity of calculations, we modeled the epidemics only among foragers without taking nurses into account. The number of healthy foragers was designated as *N*_*S*_, while the number of infected ones – as *N*_*I*_. The number of foragers infected daily was *N*_*S*_ · *N*_*I*_ · *β*, where *β* is the pathogen transmission efficiency. No recovery from infection was assumed. In this case, the fitness function becomes:

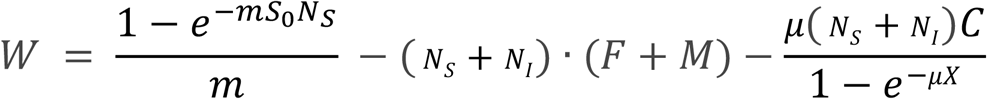

Investigation of W(X) in this model demonstrated the existence of an optimal X that defines maximum W independently of D and M. Therefore, limiting lifespan may be evolutionarily advantageous not because it allows for saving energy, but because it helps to control pathogen spread. At lower values of *X*, the pathogen may not be feasible, while at higher levels, it can propagate and even destroy the colony (Fig 3, **Supplementary Material III)**.

**Fig. 3.**
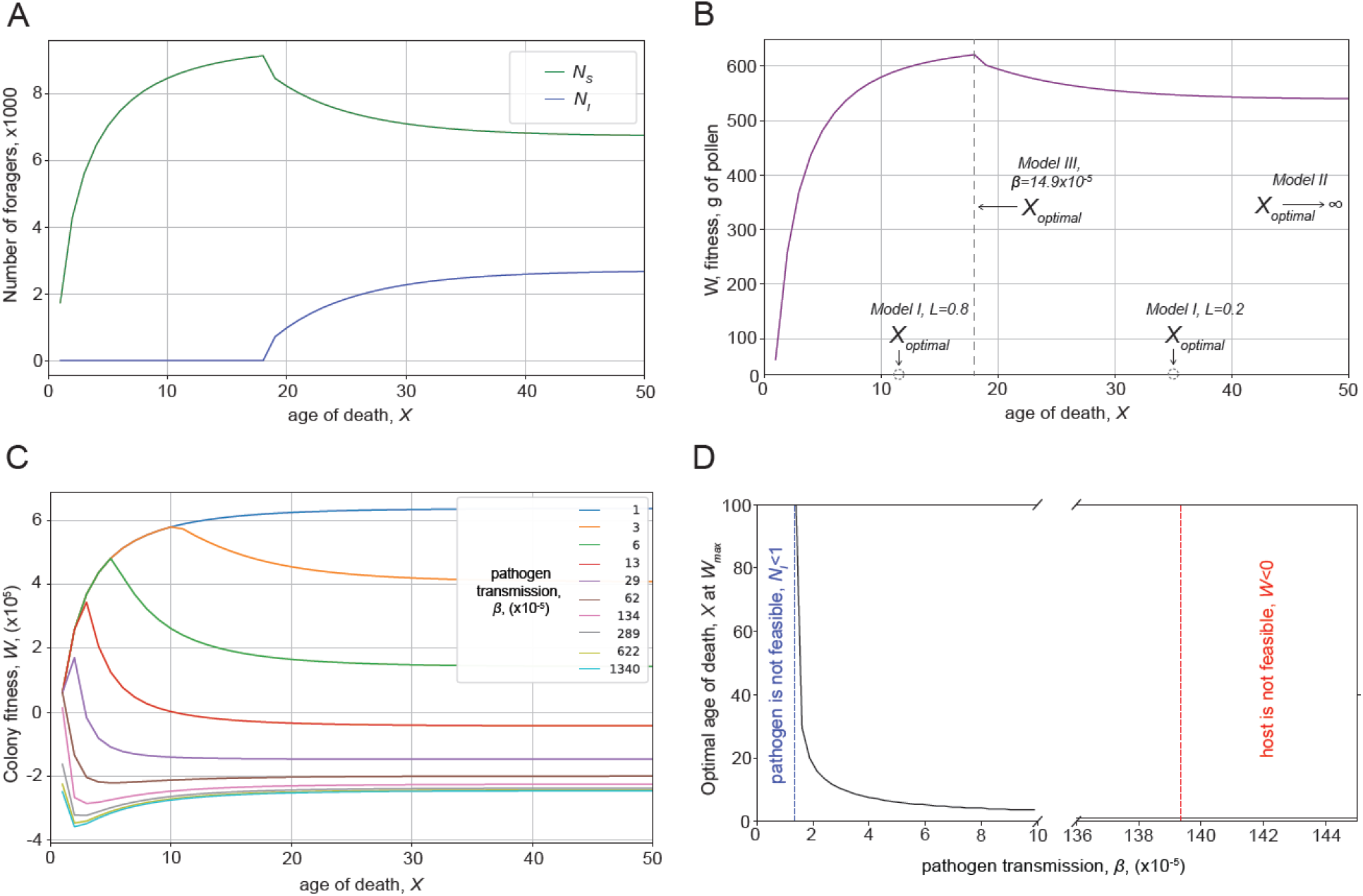
Chronic pathogens with strong effects on fitness create selective pressure towards limited lifespan. **A**. Forager’s lifespan defines the progression of epidemics. Number of infected (*N*_*I*_) and healthy (*N*_*S*_) foragers are plotted at different values of maximum lifespan *X*. Pathogen transmission is *β* = 14.9 · 10^−5^. Age-independent mortality μ = 0.139. **B**. Colony fitness (*W*) dependence on maximum lifespan *X*. Optimal parameters of *X* for all three models are shown: if *M* cannot change with time (Model I), the optimal lifespan is defined by *L* and *μ*. Although, if *M* is variable (Model II) the optimal *X* is predicted to be infinite, and aging is absent. However, epidemics may create selective pressure defining optimal lifespan (Model III). *β* and *μ* are the same as in panel A. **C**. Colony fitness dependence on maximum lifespan *X* has pronounced optimum at different values of *β*, thus justifying the evolutionary rationale of a forager’s limited lifespan. **D**. Optimal age of death dependence on pathogen transmission *β*. The optimal age of death *X* is calculated as a value under which colony fitness *W* reaches its maximum.

Therefore, limiting lifespan under the pathogen pressure may be an evolutionarily advantageous strategy. Moreover, even if we assume that damage *D* is not sufficient to mediate honeybee death at the optimal age of *X* with *M* = 0, the model formally predicts the evolution of an active killing mechanism that would increase damage and accelerate aging to ensure the ecologically optimal value of *X*. Thus, the presence of chronic debilitating pathogens may favor the evolution of earlier death and the emergence of self-damaging mechanisms for its realization. On the contrary, the activation of rejuvenation or other lifespan-prolonging adaptations may pose epidemiological risks and must be avoided.

## Discussion

Recent advances in cellular and tissue rejuvenation are remarkable and exciting. However, here we demonstrated that an optimistic interpretation of these experiments contradicts the classic theories of aging.

Selection shadow and damage accumulation consider aging as a detrimental effect that affects fitness too little to be removed by evolution. The so-called “buy now, pay later” models, antagonistic pleiotropy, and disposable soma view aging as an adverse side effect of some positive traits that improve reproduction in the youth(Gems & Kern, 2024; Kirkwood, 1977; Williams, 1957). The advantages an organism receives at the expense of aging could be increased fecundity (irrelevant for sterile workers in eusocial species) or energy saving (see **Models I** and **II**).

While these hypotheses are remarkably different, they converge on the point that aging is detrimental to fitness and predict that if an animal can rejuvenate or pause aging, utilizing this opportunity must be evolutionarily advantageous. Why, then, does the ability to reverse or pause aging not evolve ubiquitously? The classic answer would be that rejuvenation may be a complex or energetically expensive mechanism, and there is insufficient selective pressure to develop it, e.g., because cases where an old animal accesses a rich food source are too scarce. Indeed, if we accept the premise, we can explain the scarcity of rejuvenation.

However, eusocial insects pose a fundamental challenge to this logic. Workers in an eusocial community are analogous to the somatic tissue in a multicellular organism. Workers’ contribution to reproductive fitness is directly linked to the tasks they perform, rather than investment in childbirth, and as such, what can be the benefit acquired according to the “buy now, pay later” model? Additionally, members of the eusocial community have constant access to an abundant food source. Why doesn’t evolution favor long-lived workers? Is it impossible to make them more durable?

Observations suggest the opposite: aging is exceptionally plastic in eusocial animals. Queens in many species live 10 to 50 times longer than workers. While the rejuvenation exhibited in honeybees is remarkable, it remains unclear whether it is comprehensive or partial, and to what extent it increases the lifespan. Nevertheless, data from other eusocial insects indicate that workers’ lifespan can be extended enormously: the worker to gamergate transition in ants *Harpegnathos saltator* increases the lifespan ~4 times (Ghaninia et al., 2017). Tapeworm infection in ants, *Temnothorax nylanderi*, was estimated to increase lifespan more than ten times (Beros et al., 2021). If lifespan-prolonging mechanisms were already invented, why does evolution utilize them only in cases of stress applied to the colony? Let’s examine the remaining argument that extending workers’ lifespan is energetically disadvantageous.

If a worker dies, the colony must produce a new worker to replace the old one. Could this be an energy-saving strategy? During development, a honeybee larva consumes ~160 mg of pollen, while the forager’s daily ration is ~4 mg (Brodschneider & Crailsheim, 2010; Crailsheim et al., 1992). Could the cost of the worker rejuvenation or pausing aging exceed 160 mg of pollen? It does not seem possible just due to the adult honeybee’s digestive capacity. The metabolic levels of the long-lived, parasitized ants do not indicate high energy costs associated with longevity (Beros et al., 2021). Moreover, such high costs of rejuvenation contradict the laws of conservation, as replacing a proportion of damaged cells and proteins should always be cheaper than creating a new organism from scratch. Of course, the introduction of some non-additive factors that make rejuvenation unreasonably expensive may rescue the Model II. However, there are no empirical or theoretical indications that rejuvenation or other forms of lifespan elongation can be so costly. Therefore, Model II demonstrated that the reasons for the evolutionary avoidance of rejuvenation cannot be attributed to energy investment; hence, the classic “buy now, pay later” framework fails to explain the phenomenon.

It should be emphasized that our modeling concerned non-reproductive members of the eusocial colony. This social structure, in many instances, benefits the self-sacrifice of non-breeders (Joiner et al., 2016), who are functionally equivalent to somatic tissue and can be disposed of, much like skin or gut cells in the mammalian body. However, according to our calculations, even such dispensable animals must extend their lifespan if it is mechanistically possible. The effect in solitary animals is expected to be even stronger.

Finally, with **Model III**, we found that the pathogen control hypothesis provides a good fit to the observed phenomena. An epidemic of a chronic pathogen may impose selective pressure that favors shorter lifespans in foragers. As the probability of being infected increases with age, the death of aging may help remove older and potentially infected individuals. Rejuvenation or pausing aging in this model may exacerbate epidemics and should be avoided unless it is necessary for overcoming other environmental challenges, such as the death of queens, nurses, or suboptimal temperature.

Moreover, if biochemical damage received by an animal is insufficient to kill it by the optimal time, the model predicts the evolution of specialized killer mechanisms that ensure optimal lifespan. In this case, programmed aging is mathematically predicted by our model.

The pathogen control hypothesis posits aging as a programmed adaptation that evolved to control epidemics. Consistently, some chronic pathogens are expected to inhibit aging, thereby extending the host’s lifespan and infectious period. Such parasites of eusocial insects were mentioned above (Beani et al., 2021; Beros et al., 2021). Other parasites have been shown to reduce mortality and prolong lifespan in solitary insects (Hurd et al., 2001), fish (Lima et al., 2007), molluscs (Minchella et al., 1985), and even humans (Harnett & Harnett, 2024). However, the extent is much smaller than the tapeworm-mediated lifespan extension in ants *T. nylanderi* (Beros et al., 2021). It is plausible that pathogens do not introduce entirely new host lifespan-prolonging functions, but rather hijack the intrinsic aging plasticity mechanisms. If so, the presence of such mechanisms in an animal may render it vulnerable to a potential breach in pathogen defense. Hence, aging plasticity mechanisms, including rejuvenation, should not evolve unless they provide substantial selective advantages as in eusocial species. Moreover, if such mechanisms eventually become redundant, they must be eliminated by evolution to remove a potential weakness in anti-pathogen defense. This logic may explain why rejuvenation is scarce, despite being mechanistically possible.

The progress in developing rejuvenation therapies requires a solid theoretical basis. Here, we formally demonstrate that neutral aging (damage accumulation and selection shadow) and “buy now, pay later” (antagonistic pleiotropy and disposable soma) models of aging imply that rejuvenation is mechanistically impossible. Therefore, approaches striving to achieve rejuvenation based on these models contain internal contradictions. Alternative models, such as the pathogen control hypothesis (Lidsky & Andino, 2020, 2022; Lidsky et al., 2022), are required to reconcile evolutionary theory with rejuvenation. Detailed investigations of natural examples of rejuvenation are essential not only to inform the field about its feasibility, but even more importantly, to advance and refine the evolutionary theory of aging.

