## Supplementary Materials for "Avoidance Of Rejuvenation: A Stress Test For Evolutionary Theories Of Aging"

### Supplementary material I.

**Model I. Investment in maintenance is constant.**

$t$ – age of the forager

$\mu$ – age-independent mortality per day; exp(-μt) - survival as a function of age t

$n_0$ – quantity of new foragers born per day

$n(t)$ – quantity of foragers of age $t$; $n(t) = n_0 \cdot exp(-\mu t)$

$N$ – total number of foragers within the colony

C – energy needed to produce one forager, e.g., required to feed the larva.

$F$ – energy that a forager consumes daily to do its work

$D$ – the daily damage a forager receives, expressed in the energy equivalent;

$M$ – energy investment into repairing the damages; $(D - M)$ - daily unrepaired damage

$L$ – the proportion of molecules sufficient to cause death if damaged; if unrepaired damage exceeds $C \cdot L$, the forager dies.

$X$ – maximum intrinsic lifespan, days

(1) $$X = \frac{L \cdot C}{D - M}$$

(2) $$N = \int_0^X n(t)dt = n_0 \frac{1 - exp(-\mu X)}{\mu} = n_0 \frac{1 - exp(-p)}{\mu}$$

(3) $p = \mu X$ – maximum longevity, dimensionless value

$U(n)$ – production of the first $n$ foragers

$S(n)$ – production of the $n^{\text{th}}$ forager

(4) $$S(n) = \frac{dU}{dn} = S_0(1 - m \cdot U(n))$$

$\frac{1}{m}$ – maximum production of all foragers under any $n$

$W$ – reproduction fitness

(5) $$W = U(N) - N(M + F) - n_0 C$$

There must be optimal $n_0$ and $M$ resulting in maximum $W$.

(6) $$U(n) = \frac{1 - exp(-m \cdot S_0 \cdot n)}{m}$$

(7) $$S(n) = S_0 \cdot \exp(-m \cdot S_0 \cdot n)$$

(8) $$n_0 = \frac{\mu N}{1 - exp(-p)}$$

Therefore:

(9) $$W = \frac{1 - exp(-m \cdot S_0 \cdot N)}{m} - N(M + F) - N \frac{\mu C}{1 - exp(-p)}$$

(10) $$\frac{dW}{dN} = S_0 \cdot \exp(-mS_0 n) - (M + F) - \frac{\mu C}{1 - \exp(-p)}$$

As $\frac{dW}{dN} = 0$, there is an equation

(11) $$S_0 \cdot exp(-mS_0 N) = M + F + \frac{\mu C}{1 - exp(-p)}$$

Let us declare dimensionless parameters: $\lambda = \frac{M + F}{S_0}$; $\gamma = \frac{\mu C}{S_0}$;

Therefore $N = -\frac{1}{mS_0} \cdot ln(\lambda + \frac{\gamma}{1 - exp(-p)})$

Let assign η = $\lambda + \frac{\gamma}{1-e^{-p}}$ and insert the equation for N into equation (9) that defines W.

We would receive:

(12) $$W = \frac{1}{m}(1 - \eta + \eta \cdot \ln(\eta))$$

To find optimal values warranting maximal fitness, we need to solve the equation $\frac{dW}{dM} = 0$

However, $\frac{dW}{d\eta} = \frac{ln(\eta)}{m} < 0$ . Therefore, we solve the equation $\frac{d\eta}{dM} = 0$.

(13) $$\frac{d\eta}{dM} = \frac{1}{S_0} - \frac{\gamma \cdot exp(-p)}{(1-exp(-p))^2} \cdot \frac{p}{D-M}$$

After multiplying the two parts on $\mu CL$, the equation takes the final form:

(14) $$\frac{p^2 \cdot exp(-p)}{(1-exp(-p))^2} = L$$

### Supplementary material II.

**Model II. Investment in maintenance is variable.**

The expression (1) for X now becomes:

$$(15) \quad D \cdot X - L \cdot C = \int_0^X M(t)dt$$

Now the total compensation for bees is instead of $M \cdot N$ is

$$(16) \quad \int_0^X n(t)M(t)dt$$

Therefore:

$$(17) \quad W = \frac{1-exp(-m \cdot S_0 \cdot N)}{m} - NF - \frac{\mu N}{1-exp(-p)} \int_0^X \exp(-\mu t)\, M(t)\, dt - N \frac{\mu C}{1-exp(-p)}$$

Here, we used the equation:

$$(18) \quad n(t) = N \cdot \frac{\mu \cdot exp(-\mu t)}{1 - exp(-p)}$$

M(t) must be chosen in a way that:

$\int_0^X M(t)dt$ is constant, while $\int_0^X \exp(-\mu t)M(t)dt$ must be as low as possible.

Let define:

$X_0 = \frac{L \cdot C}{D}$; so $M(t) = \begin{cases} 0, & t < X_0 \\ D, & X_0 < t < X \end{cases}$

Applying this to equation 17 brings us to:

$$(19) \qquad W = \frac{1-exp(-m \cdot S_0 \cdot N)}{m} - NF - \frac{ND(exp(-\mu x_0) - exp(-p))}{1-exp(-p)} - N\frac{\mu C}{1-exp(-p)}$$

From $\frac{dW}{dN} = 0$ it turns out:

$$(20) \qquad \exp(-mS_0N) = \frac{F}{S_0} + \frac{D}{S_0} \cdot \frac{\exp(-po)-\exp(-p)}{1-\exp(-p)} + \frac{\gamma}{1-\exp(-p)}$$

Here $p_0 = \mu X_0$

Let’s define the right part of the equation as η. Therefore:

$$(21) \qquad N = -\frac{1}{mS_0} ln(\eta)$$

Therefore:

$$(22) \qquad W = \frac{1}{m}(1 - \eta + \eta \cdot ln(\eta))$$

As $\frac{dW}{d\eta} = \frac{ln(\eta)}{m} < 0$; and

$$(23) \qquad \frac{d\eta}{dp} = \frac{exp(-p)(D(1 - exp(-\frac{\mu \cdot L \cdot C}{D})) - \mu C)}{S_0(1-exp(-p))^2} < 0$$

then $\frac{dW}{dp} > 0$, an increase in $p$ should always increase the colony fitness $W$, and death by old age should never be observed.

### Supplementary material III.

**Model III. Infections provide selective pressure for limiting lifespan.**

$N_I$ - quantity of infected foragers. Infected individuals do not contribute to foraging but consume resources as uninfected colony members ($N_S$):

$$(24) \qquad W = \frac{1-e^{-mS_0N_S}}{m} - (N_S + N_I)\cdot(F+M) - \frac{\mu(N_S+N_I)C}{1-e^{-\mu X}}$$

The probability of being infected per day is - $\beta N_I$.
The working bees are being infected or dying with the probability:

$$(25) \qquad \mu + \beta N_I = \mu(1 + \nu N_I), \text{ where } \nu = \frac{\beta}{\mu}.$$

We suppose that:

$$(26) \qquad n_0 = -\frac{\mu}{(1-exp(-p))\cdot mS_0} ln(\lambda + \frac{\gamma}{1-exp(-p)})$$

as in Model I. Therefore:

$$(27) \qquad N_S + N_I = \int_0^x \cdot\, n_0 \cdot\, exp(-\mu t)dt = n_0 \cdot \frac{1-exp(-p)}{\mu}$$

$$(28) \qquad N_S = \int_0^x n_0 \cdot exp(-\mu(1+\nu N_I)t)dt = n_0 \cdot \frac{1-exp(-p(1+\nu N_I))}{\mu(1+\nu N_I))}$$

Receiving the equation for $N_I$:

(29) $$N_I = \frac{n_0}{\mu} \cdot \frac{\nu N_I - (1+\nu N_I)exp(-p) + exp(-p(1+\nu N_I))}{1+\nu N_I}$$

This equation always has a solution $N_I = 0$

Let's assign the right part of the equation as $\varphi(N_I)$. When the values of $N_I$ are big:

(30) $$\varphi(N_I) \sim \frac{n_0}{\mu}(1 - exp(-p))$$

If $\frac{d\varphi}{dN_I} \Big|_{N_I=0} > 1$, under small $N_I$ values $N_I < \varphi(N_I)$ and the equation's solution $N_I = \varphi(N_I)$ under $N_I > 0$ exists.

However, $\frac{d\varphi}{dN_I} \Big|_{N_I=0} = 1$ could be considered as an equation on $p$. Its solution is the value of $p$ under which the infection cannot propagate, as an average forager infects less than one healthy forager per lifespan ($R_0 < 0$).

The final equation is:

(31) $$-\frac{\nu}{mS_0} \cdot \frac{1-(p+1)exp(-p)}{1-exp(-p)} \cdot ln(\lambda + \frac{\gamma}{1-exp(-p)}) = 1$$

The stationary solution exists if a constant $\frac{\nu}{mS_0} \sim 1$.
